# Bumblebees learn a relational rule but switch to a win-stay/lose-switch heuristic after extensive training

**DOI:** 10.1101/2020.05.08.085142

**Authors:** HaDi MaBouDi, Cwyn Solvi, Lars Chittka

## Abstract

Mapping animal performance in a behavioural task to underlying cognitive mechanisms and strategies is rarely straightforward, since a task may be solvable in more than one manner. Here, we show that bumblebees perform well on a concept-based visual discrimination task, but spontaneously switch from a concept-based solution to a simpler heuristic with extended training, all while continually increasing performance. Bumblebees were trained in an arena to find reward on displays with shapes of different sizes where they could not use low-level visual cues. One group of bees was rewarded at displays with bigger shapes and another group at displays with smaller shapes. Analysis of total choices shows bees increased their performance over 30 bouts to above chance. However, analyses of first and sequential choices suggest that after approximately 20 bouts, bumblebees changed to a win-stay/lose-switch strategy. Comparing bees’ behaviour to a probabilistic model based on a win-stay/lose-switch strategy further supports the idea that bees changed strategies with extensive training. Analyses of unrewarded tests indicate bumblebees learned and retained the concept of relative size even after they had already switched to a win-stay, lost-shift strategy. We propose that the reason for this strategy switching may be due to cognitive flexibility and efficiency.

## 1. Introduction

Cognitive flexibility reflects an individual’s ability to adaptively alter their behavioural strategy in accordance with a changing environment [1]. A fundamental challenge for animal cognition researchers is to decipher which strategies an animal uses in solving any particular task [2]. Indeed, there is often multiple ways for an animal to solve a behavioural task.

Bees have been shown capable of learning various abstract relationships for example rules about target size (e.g. “pick the larger (or smaller) of two object sizes”), amongst myriad impressive cognitive abilities [3–5]. However, in some of these cases it may be that bees use a variety of different strategies to solve the tasks they are confronted with [5,6]. One recent study showed that some bees can solve a spatial concept learning task using a simple visual discrimination strategy through sequential scanning of stimuli rather than needing to compare stimuli based on an abstract rule, though other individuals may well follow such a rule [7]. In some numerical cognition tasks, honeybees may also use alternative cues that correlate with number, but are not in themselves numerical [8]. Bees’ behavior in solving a delayed matching-to-sample task has been shown to be replicated by a model without any neural representations of the abstract concepts of sameness or difference [6]. Even the same individuals may have recourse to different solutions to the same task, depending on the extent of training. For example, with an increased number of training trials with a single pair of patterns, individual honeybees have been shown to have a greater generalized response to novel stimuli, i.e. the representation necessary to discriminate subsequent visual patterns changes with extended training [9]. All of these findings highlight the need for considering alternative strategies used by animals in cognitive tasks. This does not just concern the traditional dichotomy of “simple” versus “complex” solutions to such tasks. Different individuals may use different solutions that are equal in complexity, depending on their particular path to figuring out a solution.

Previous works have shown that honeybees are able to solve a task that appears to necessitate learning the concept of relative size and apply the rule to novel sizes within or outside the size range they were trained [10,11]. As with the examples above, bees may use more than one strategy to solve the same task, depending on the training protocol and context. Here, we test bumblebees to determine the strategies by which they cope with a relational rule learning task (“bigger-than”/”smaller-than”) and examine their behaviour over time to reveal the cognitive strategies used over the course of training.

## 2. Material and methods

### (a) Animals and experimental setup

Bumblebees (*Bombus terrestris audax*) from commercially available colonies (Agralan Ltd, UK), were housed in a wooden nest-box connected to a flight arena (100 cm x 75 cm x 30 cm). Bees were allowed access to a flight arena through an acrylic corridor (25 cm x 3.5 cm x 3.5 cm). Three plastic sliding doors located along the corridor allowed controlled access to the arena. The arena was covered with a UV-transparent clear acrylic sheet. The stimuli were presented to bees on the grey-coloured back wall of the arena.

Although there are no current requirements regarding insect care and use in research, experimental design and procedures were guided by the 3Rs principles [12]. The bumblebees were cared for on a daily basis by trained and competent staff, which included routine monitoring of welfare and provision of correct and adequate food during the experimental period. Colonies were provided with ∼7 g irradiated commercial pollen (Koppert B.V., The Netherlands) every two days. Bees from three colonies were used in this study.

### (b) Pretraining phase

All bumblebee workers were recruited from a gravity feeder containing 30% (w/w) sucrose solution placed in the centre of the arena. Outside of experiments, the colony was provided with 30% (w/w) sucrose solution from a small gravity feeder placed inside the nest-box during the evenings. Successful foragers on the arena gravity feeder were individually marked with number tags, superglued to their thorax, for identification during the subsequent experiment (Opalithplättchen, Warnholz & Bienenvoigt, Ellerau, Germany). Marked bees were pre-trained to find 50% (w/w) sucrose solution from microcentrifuge tubes (5mm diameter) at the centre of each of six white discs (7 cm diameter) on the grey-coloured back wall of the arena, horizontally 14 cm from each other vertically 9.3 cm from each other (positioned as in figure 1). These discs were made of paper and covered with transparent laminate to enable cleaning with 70% ethanol in water (v/v). All stimuli were printed with a high-resolution printer.

**Figure 1.**
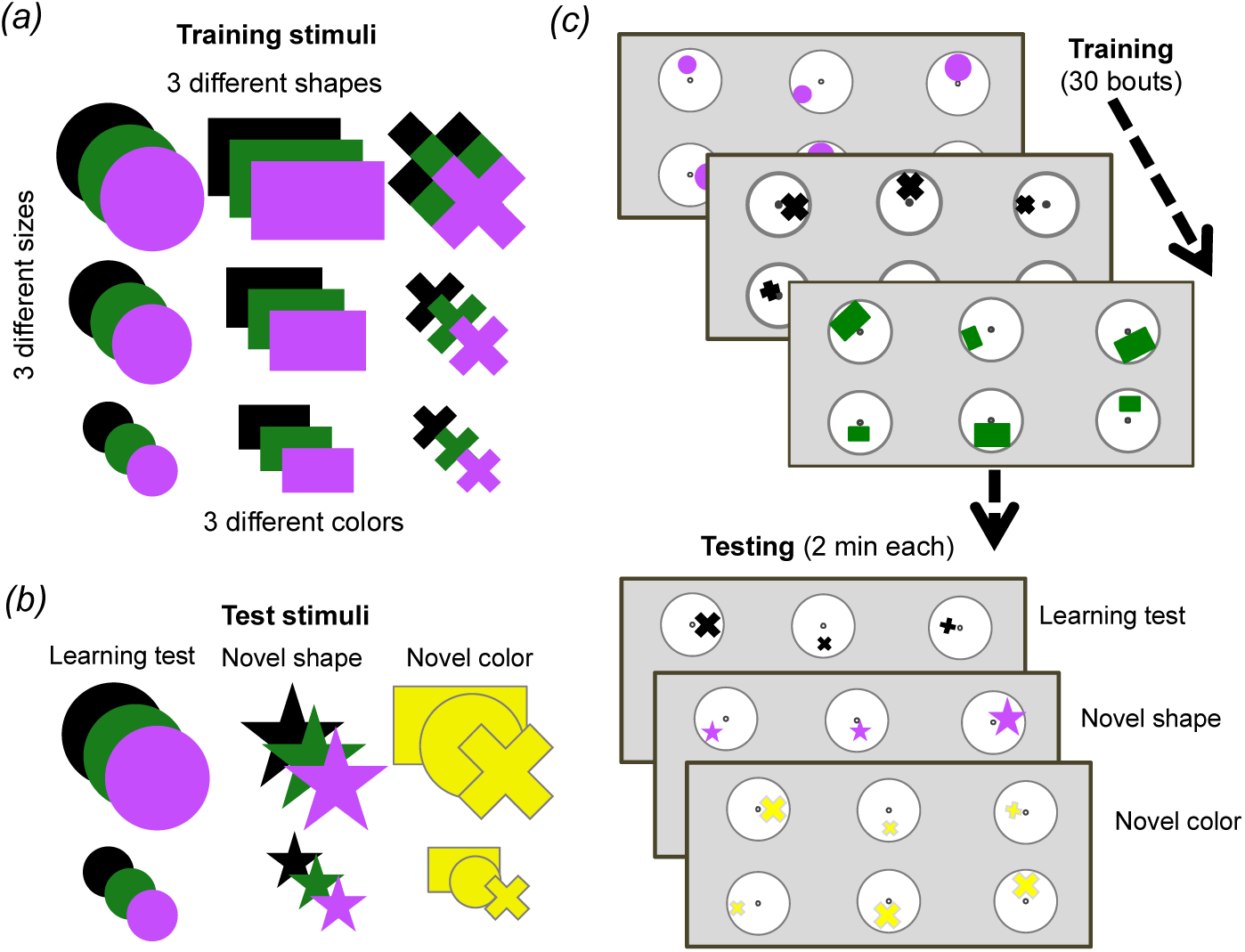
Training and testing protocol. *(a)* Stimuli options used during training. *(b)* Stimuli options used for each of the three different unrewarded tests. *(c)* Training and test protocol. Bees were trained for 30 bouts (visits to the arena before returning to the hive). All stimuli in (*a*) were used randomly across bouts during training. Only two of the possible three sizes of shapes were presented during a single bout. Only one of the possible three colours and one of the possible three shapes were presented each bout. One group of bees (n = 10) was trained to find 50% sucrose solution at the centre of the stimulus containing the bigger of the three shapes and bitter quinine solution at the smaller of the three shapes. Another group (n = 8) were trained on the opposite contingency. Once training was complete, bees were subjected to three unrewarded tests (with one or two reminder/training bouts between each test to keep bees motivated). All tests used small and large sized shapes. The learning test used one randomly chosen type and colour used during training. The novel shape test used one randomly chosen colour used during training but always a star shape that had not been used during training. The novel colour test used one randomly chosen shape used during training but always coloured yellow, which had not been used during training.

### (c) Training phase

When several number-tagged bees had learnt to find reward from the tubes located in the centre of the display discs, one bee was randomly selected for the training phase. During the training phase, an individual bee was trained on six discs on the back wall of the arena with the same spacing as in training, each displaying one of two different sized shapes (figure 1*a* and *c*). During 30 training bouts (a bout was a bee’s visit to the arena and landings on different stimuli until satiated and subsequent return to her nest) one group of bees (n = 10) learned that the bigger of the two shapes contained 30 *µ*l 50% sucrose solution and the smaller contained 30*µ*l saturated quinine hemisulfate solution (bigger-than rule). Another group of bees (n = 8) learned the reverse contingency (smaller-than rule).

Between training bouts, each disc was rotated randomly so that the position of a shape varied across the six discs in relation to the central microcentrifuge tube containing sucrose solution (figure 1*c*). The location, shape and colour of stimuli sets were changed between bouts. The shapes used in training varied in size (small, medium, large), type (circle, rectangle, cross) and colour (black, green, purple) (figure 1*a*). Only one type and colour of stimuli was presented to a bee in each bout and only two of the three sizes were presented during one bout. The dimensions of the shapes were as follows: small circle: Ø = 1.07 cm; medium circle: Ø = 1.97 cm; large circle: Ø = 2.87 cm; small rectangle: 0.93 cm x 1.18 cm; medium rectangle: 1.79 cm x 2.92 cm; large rectangle: 2.3 cm x 3.94 cm; small cross: width of bars = 0.46 cm, length of bars = 1.3 cm; medium cross: width of bars = 0.6 cm, length of bars = 2.15 cm; large cross: width of bars = 0.96 cm, length of bars = 2.87 cm. Note that there was a large variability between physical features of stimuli (figure *S1*). This variability ensured the bees were not able to solve the task by associating an absolute size of stimuli with certain reinforcements. Several of stimuli were paired with both positive and negative reinforcements during the training phase. For instance, middle size stimuli were paired with the positive reinforcement in some training bouts while these were paired with negative reinforcement in the rest of the training bouts. Further, the total area of the middle rectangular was bigger than the total area of the big cross (see figure S1a). All of these variations described ensured that low-level visual cues could not be used to solve the task. Stimuli were cleaned between each training bout with 70% ethanol in water (v/v) to ensure odour cues were not used to solve the task. After the daily experiment, all used microcentrifuge tubes were washed with soap-water, then cleaned with 70% ethanol solution. Finally, they were rinsed with water and air-dried in the temperature of the lab during the night.

### (d) Testing phase

Following the training phase, each bee was tested in the same setup as in training in three different scenarios, but with stimuli in the tests providing 30 μl of sterilized water (figure 1*b and c*). Tests lasted 120 seconds, at which point the bee was gently removed from the arena by using a cup and placed into the corridor until stimuli were changed for the refreshment bouts. Each test was separated by two refreshment training bouts between tests to maintain the bee’s motivation.

The sequence of the three tests were counterbalanced across bees. The learning test evaluated performance by testing bees on one of the same sets of stimuli used during training, pseudo-randomly chosen (i.e. a random number generator was used to generate a random sequence of tests for each individual bee). The learning test used only the small- and bigger-sized training shapes. The other two tests used either a novel shape and size (star) or a novel colour (yellow), with the other properties pseudo-randomly chosen. The dimensions of the 5-poited stars were as follows: small star: length of side of point = 0.5 cm; large star: length of side of point = 1.23 cm (See figure S1). As in training, stimuli were cleaned between each bout during the testing phase with 70% ethanol in water (v/v) to ensure odour cues were not used. Trained bees were removed from the nest once the training and tests phases were finished.

### (e) Statistical analysis and probabilistic model of learning curve

To evaluate bees’ performance over bouts, the percentage of correct choices (choices were defined as when a bee touched a microcentrifuge tube with her antennae or when she landed on a microcentrifuge tube) was calculated from either all choices or from only the first or second choices within each block of six bouts during training (total of five blocks). Using a generalised linear mixed model (GLMM) for binary probability (correct or incorrect), the effect of different factors such as colony, group of training and interaction between trial block and group of bees in the bees’ performance were calculated. The bee index was included in the model as random factors. GLMMs were performed in MATLAB (MathWorks, Natick, MA, USA).

To determine whether bees used relative size information, rather than any other visual cues, the choices of bees during the unrewarded tests were evaluated by a Wilcoxon signed rank test. Further, a Kruskal-Wallis test was used to statistically evaluate and compare whether the bees’ performance or choice numbers in different blocks of bouts are from the same distribution.

To test if bees might use a win-stay/lose-switch strategy during training, we calculated the conditional probabilities of each bee’s second choice (*c*_2_) given their first choice (*c*_1_) at each block of 10 bouts. A conditional probability, “Probability of B, given A (*P*{*B*|*A*})”, is a probability of an event (B) occurring given that another event (A) has already occurred. The conditional probability of a lose-switch strategy, i.e. a correct second choice after an incorrect first choice, is calculated by *P*{*c*_2_= 1|*c*_1_= 0} = P{*c*_2_= 1,*c*_1_= 0}/*P*{*c*_1_= 0} where *P*{*c*_2_= 1,*c*_1_= 0} is the joint probability of a correct second choice and an incorrect first choice and *P*{*c*_1_= 0} is the probability of the first incorrect choice. The conditional probability *P*{*c*_2_ = 1|*c*_1_ = 0} at more than chance level indicates that a bee switched to another presented size when they found the first choice was incorrect. In the same way, we can calculate the conditional probability of a win-stay strategy, using *P*{*c*_2_ = 1|*c*_1_ = 1} = *P*{*c*_2_ = 1, *c*_1_ = 1}/*P*{*c*_1_ = 1}, i.e. the bee’s second choice is the same size as the first choice when their first choice was correct.

#### Model of prediction of learning curve based on a bee’s first two choices

We propose a Markov stochastic model [13] to describe the learning curve of bees’ choices (total choices at each bout) based on the information of two first choices of bees. The performance of the model at each bout is assumed as 

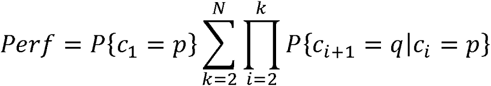

*P*{*c*_1_} is the probability of the first choice at each bout and *P*{*c*_*i*+1_| *c*_*i*_} is the conditional probability of (*i* + 1) − *th* choices given the of *i* − *th* choices (*i* ≥ 1) for when each choice in the sequential choices is correct or incorrect. *p or q* = 1 if the choices are correct, otherwise *p or q* = 0. We assume that the conditional probabilities of two sequential choices from the third choices are equal to the conditional probability of the second choice given the first choice expressed by bees at each bout of training. The sequence of possible events in which the probability of each event depends only on the state achieved in the previous event will be stopped (N) when the simulated bees collect all three positive reinforcements along with two, one or no incorrect choices within each bout according to the average number of choices at each bout.

## 3. Results

### (a) Bees’ overall performance increased over the 30 training bouts

A multivariate statistical model, GLMM, applied to the performance of bees demonstrates a significant increase in the proportion of correct choices made over the 180 choices of the training phase (figure 2*a*, p = 0.018) irrespective of the shape, colour or position of patterns within the stimuli. No significant differences were found between the learning curves of the two different contingency groups (*i.e.* “bigger-than” rule versus “smaller-than” rule; p = 0.87). The output of the GLM confirms that there was no significant difference between the different colonies of bees during the training phase (p = 0.37). These results show that bees became better at solving either contingency over training bouts.

**Figure 2.**
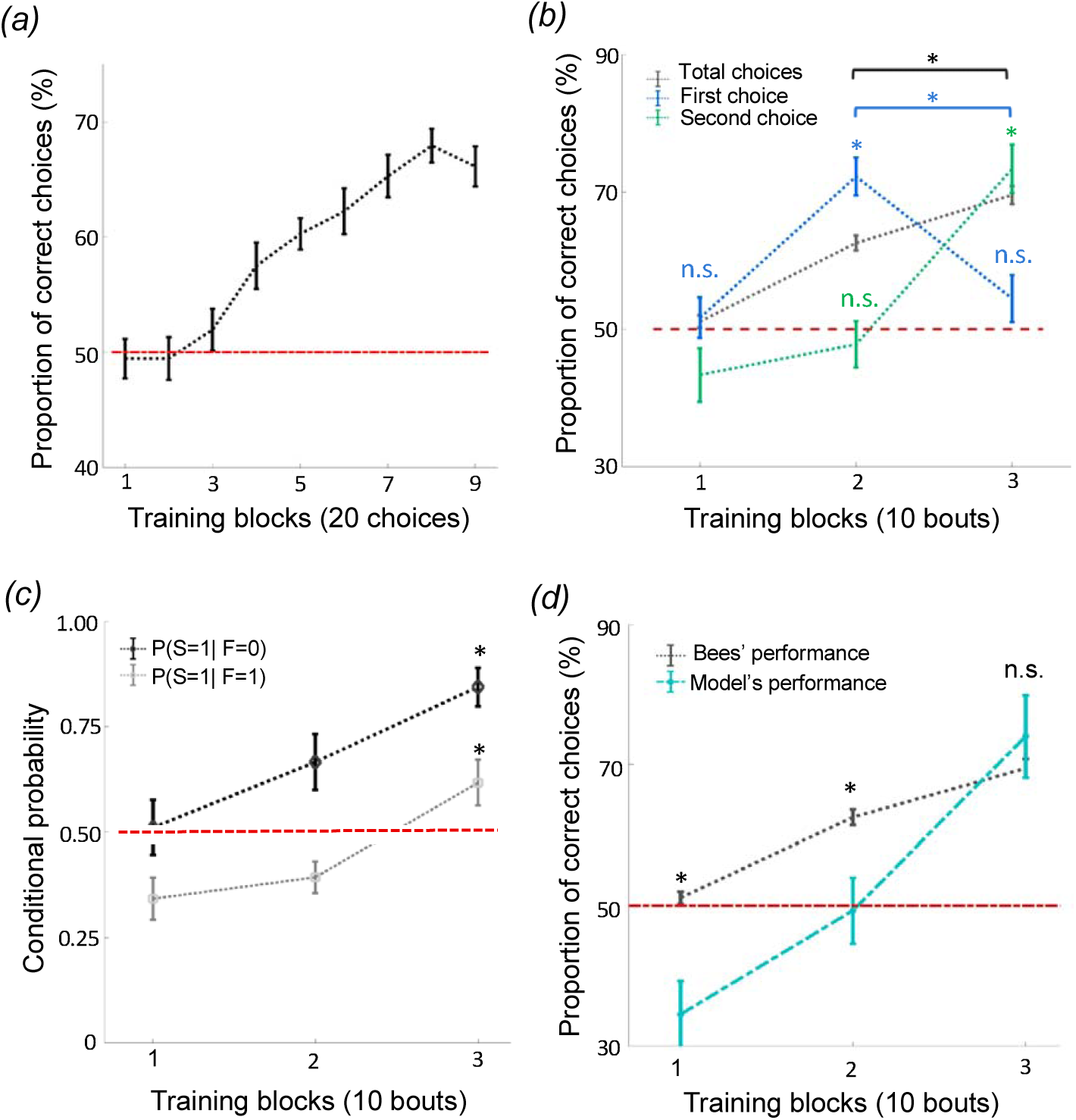
Bees use a win-stay/lose-switch strategy after extensive training. *(a)* There was a significant increase in the number of correct choices over the 180 conditioned choices (p = 0.018). *(b)* Bees’ performance over 3 blocks of 10 training bouts during the relative size discrimination task. Performance increased gradually over bouts when considering total number of choices in each bout (black line; p = 1.96e-4). Bees’ first choice performance increased significantly from the first to the second block of training bouts to 72.22% (p = 3.71e-4) but then dropped to chance level from the second to third block of training bouts (blue line; p = 0.79). Second choice performance was near chance for the first two blocks of training bouts (p > 0.49), but then increased significantly during the third block of training bouts (green line; p = 4.28e-4). These results indicate that bees changed to a win-stay/lose-switch strategy after extensive training. Vertical lines = standard error of the mean. Red dashed line = chance level performance (50%). *(c)* The average conditional probabilities of a bee’s second choice within each bout being correct given the outcome of the bee’s first choice of the bout (either correct or incorrect). Both conditional probabilities increased to above chance during the second and third blocks of bouts (p = 0.02 for win-stay and p = 2.75e-4 for lose-switch) *(d)* Our win-stay/lose-switch model’s performance matches our bees’ performance on the task during the last block of 10 bouts during training (p=0.15), again suggesting that after extensive training bees changed to a win-stay/lose-switch strategy. (Vertical lines = standard error of the mean). Red dashed line = chance level (50%).

### (b) Bees used a win-stay/lose-switch strategy after extensive training

The typical analysis used to determine whether an animal has solved a particular task is to calculate the animal’s performance based on the number of correct and incorrect choices throughout the training phase. At first inspection, bees’ behaviour during training suggests they learned to solve the concept-based task (figure 2*a*). However, a finer examination of their choices suggests the involvement of another strategy in the later stages of training. If bees had only used the concept of relative size throughout training, their first choices should reflect this by increasing in accuracy throughout the 30 bouts. Although bees’ average overall accuracy gradually increased to 70% (significantly above chance level) over the 30 training bouts (figure 2*b*; Wilcoxon signed rank test: z = 3.72, n = 18, p = 1.96e-4), their first-choice accuracy rose to 72% (significantly above chance level: Wilcoxon signed rank test: z = 3.55, n = 18, p = 3.71e-4) over the first 20 bouts and then decreased to chance level (54%) over the next 10 bouts (Wilcoxon signed rank test: n = 18, z =1.25, n = 18, p = 0.21; figure 2*b*) and decreased significantly across last two blocks of bouts (Wilcoxon signed rank test: z = 2.83, n = 18, p = 4.59e-3). Second-choice accuracy was not different from chance level during the first two-thirds of the training phase (Wilcoxon signed rank test: z = -0.67, n = 18, p = 0.49), but increased in the final third of the training phase to 73.33%, significantly above chance level (figure 2*b*; Wilcoxon signed rank test: z = 3.52, n = 18, p = 4.28e-4). These results suggest that bees changed to a win-stay/lose-switch strategy after around 20 bouts of training, i.e. if they find reward at a stimulus they choose the same type of stimulus next, or if no reward is found at a stimulus they choose a different type of stimulus next.

To help evaluate the possibility that bees switched strategies part way through training, we calculated the conditional probabilities (Materials and Methods) for 1) a correct second choice after a correct first choice (win-stay), and 2) a correct second choice after an incorrect first choice (lose-switch). Both of these two conditional probabilities increased over bouts (figure 2*c*; Kruskal-Wallis test, chi-sq > 12.94, df = 53, p < 1.55e-3), most notably rising to significantly above chance level in the last third of training (Wilcoxon signed rank test: z = 2.18, n = 18, p = 0.02 for win-stay and z = 3.63, n = 18, p = 2.74e-4 for lose-switch), again suggesting that bees had changed to a win-stay/lose-switch strategy.

### (c) Modelling a win-stay/lose-switch strategy

To further examine whether bees switched strategies during training, we utilised a probabilistic model based on a win-stay/lose-switch strategy. Within our model, we used bees’ overall and conditional performance (figures 2*b* and 2*d*) and initial first and second choices to predict bees’ subsequent choices in each bout (Materials and Methods). Figure 2*d* shows that our model predicts the bees’ performance in the last 10 bouts (i.e. no difference between the model’s performance and bee’s performance; Wilcoxon signed rank test: z = -1.41, n = 18, p = 0.15). In contrast, our model’s predicted performance was significantly poorer than the performance of bees in the first 20 bouts (Wilcoxon signed rank test: z > 2.32, n = 18, p < 0.01 for both first two blocks). The ability of our model to predict the behaviour of our bees in the later stages of training but not the initial stages supports the hypothesis that bees changed to a win-stay/lose-switch strategy within the last 10 bouts of training.

### (d) Bees retained the concept of relative size after having switched strategies

So far, our analyses and model results suggest that bees used a win-stay/lose-switch strategy only after extensive training. Bees seemed to have used a different strategy during the initial blocks of training bouts. Their increased performance to above chance level, suggests they were discriminating the stimuli based on size. To ensure that bees’ initial strategy had actually been a relative size rule, we measured bees’ performance directly after training in unrewarded tests. Because the tests were unrewarded, bees could not solve the task based on a win-stay/lose-switch strategy. Bees’ performance on the learning test was above chance level (Wilcoxon signed rank test: z = 3.73, n = 18, p = 1.87e-4), as was their performance on the novel shape transfer test (Wilcoxon signed rank test: z = 3.51, n = 18, p = 4.46e-4), and on the novel colour transfer test (z = 3.03, n = 18, p = 2.41e-3 for novel colour; figure 3*a*). Note that the variability in different shape sizes and resulting overlap between sizes across shapes prevented bees from associating a general size with reward (figure *S1*). These results suggest that the bees had at some point during training learned to solve the task based on the concept of relative size.

**Figure 3.**
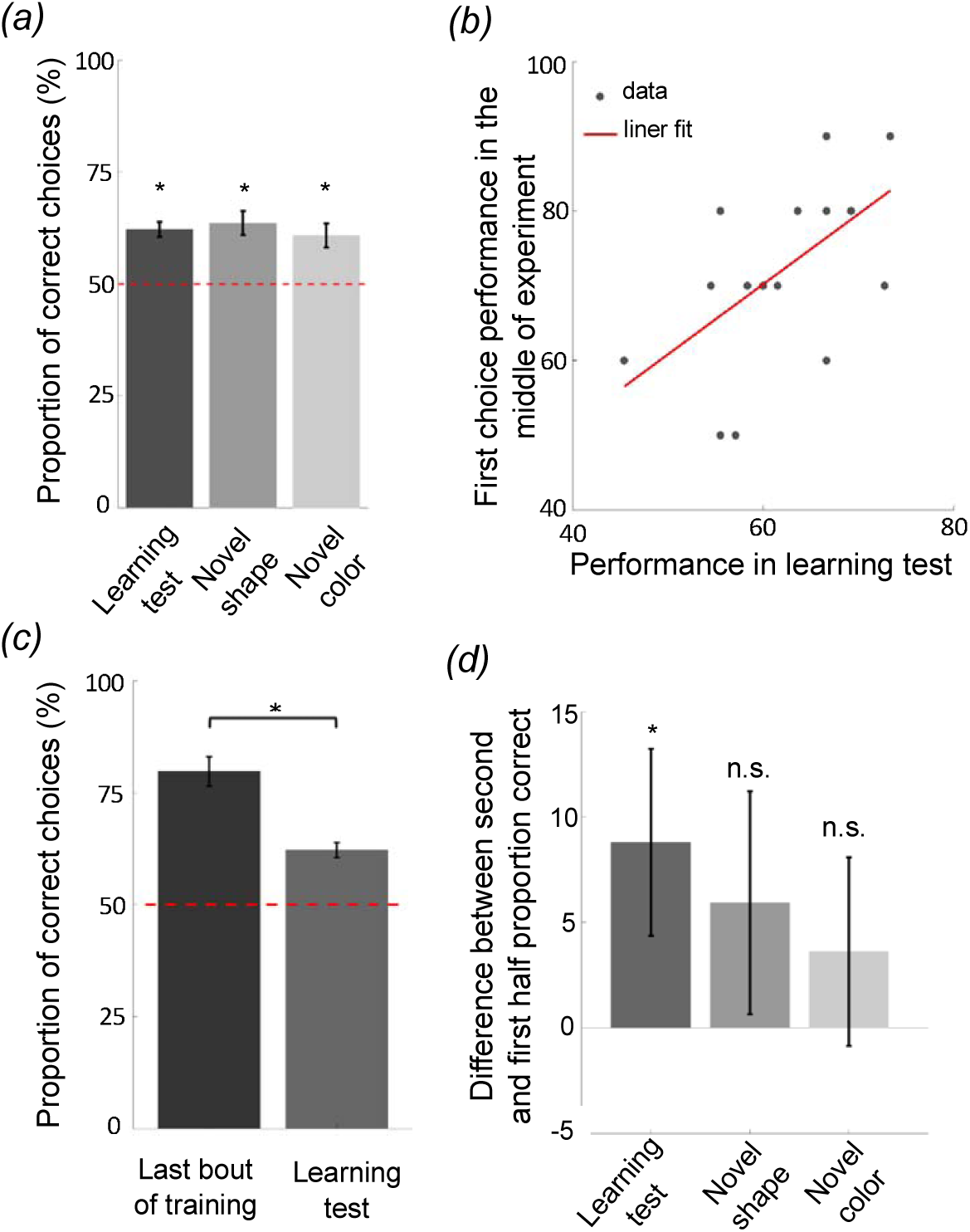
Bees learn and retain a relative size rule. *(a)* The performance of bees during each of the three unrewarded tests shows that they learned and retained the concept of relative size (p <2.41e-3). *(b)* The scatter plot displays the correlation between the performance of bees in the learning test and their first choice performance prior to changing strategies, during the second block of 10 bouts (rho = 0.58, p = 0.01). The red solid line = line of best fit. (*c and d*) The significant drop in performance from the last bout of training to the learning test (p=9.30e-4) (*d*) and the difference in performance between the second and first half of choices during each of the tests (p = 0.03 for learning test; p = 0.28 for novel shape transfer test; p = 0.14 for novel colour transfer test) suggest that bees had begun the tests with the win-stay/lose-switch strategy. Bars = mean. Vertical lines = standard error of the mean. Red dashed line = chance level (50%).

Because animals vary in their learning and performance, we posited that if bees had learned and retained a relative size rule, how well they performed in training before changing strategies should reflect how well they perform (i.e. remember the relational rule) during the learning test. In line with this, there was a positive correlation between the average of first choice accuracy in the second third of the training phase (prior to strategy change) and bees’ performance in the learning test (figure 3*b*; Spearman correlation: rho = 0.58, n = 18, p = 0.01). Although bees seemed to have changed strategies after extensive training, the results of the unrewarded tests show that bees had learnt the relative size rule during training, retained the rule even after having changed strategies late in training, and therefore resorted to the relative size rule strategy during the tests.

Note that the performance of bees in the learning test was significantly poorer than the last bout of the training phase (figure 3*c*; Wilcoxon signed rank test: z = 3.31, n = 18, p = 9.30e-4). This suggests that bees began the learning test using a win-stay/lose-switch strategy. This makes sense because they had just been using a win-stay/lost-switch strategy during training and had no knowledge that the test was unrewarded. Further, bees’ performances on the second half of choices during each of the tests was better than their performance on the first half, (figure 3*d*; Wilcoxon signed rank test: z = 1.82, n = 18, p = 0.03 for Learning test; z = 0.57, n = 18, p = 0.28 for Novel shape; z = 1.05, n = 18, p = 0.14 for Novel colour), indicating that bees had reverted to the retained relative size strategy.

Why would bees change strategies if they were already performing above chance level? We hypothesized that bees might change strategies if the new strategy was more efficient, i.e. it took them less effort to locate all three rewarding discs (discs were not refilled during training). In support of this, the number of total choices by bees decreased from an average of 7.1 choices per bout at the beginning of training to an average of 5.1 choices per bout at the end of training (figure 4; Kruskal-Wallis test, chi-sq = 22.70, df = 53, p = 1.17e-5), indicating that bees’ efficiency increased during training across a change in strategy.

**Figure 4.**
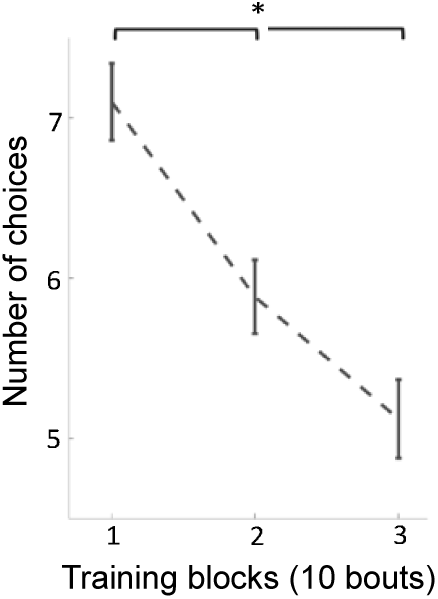
The average of number of choices on stimuli (correct and incorrect) over 3 blocks of 10 training bouts. Over training bouts, bees made fewer choices to visit all three available rewarding stimuli (p = 1.17e-5), indicating that bees continually increased efficiency on solving the task during training. Vertical lines = standard error of the mean.

## 4. Discussion

We demonstrate and corroborate previous findings [11,14] that bees can learn a relative size rule, but after extensive training opted to use a simpler strategy. Because there can often be more than one way of processing the same stimuli to solve a cognitive task, it is useful to examine individual strategies and over extended periods to explore if multiple strategies might be at play. In our paradigm, we prevented bees from using low-level visual cues. Initial increases in performance suggested that bees learned the task and later performance on unrewarded tests verified that bees had learnt and retained a relational rule, as was previously demonstrated in honeybees [11,14]. However, statistical analyses showed that after extensive training, bees began to use a win-stay/lose-switch strategy based on whether or not they were rewarded on each stimulus. Bees’ performance calculated by their first choices or by multiple sequential choices revealed a strategy of decision making that had been hidden within the gross calculation by total choices. Averaging all choices in a training bout or test is common within bee cognition and within other animal research communities. We suggest that interpretations of any animal cognition study involving multiple choices include analyses of first and sequential choices to investigate potential alternative strategies.

Theoretical and empirical work maintain that animals tend to follow the “law of least effort” [15,16], whereby subjects choose strategies that minimise the costs in obtaining desirable outcomes [15,17]. In comparative cognition research, animals may use strategies different to those we intend a specific paradigm to test and still perform well on the behaviour we are measuring [2,7,8,18,19]. Most studies have focused on the idea that animals opt to minimize physical work, but this idea extends to cognitive effort as well [20–22]. The ability to change decision-making strategies with the changing demands of the environment is essential to adaptive behaviour, and therefore survival. Lloyd and Dayan [23] proposed that constant monitoring of information to promptly assess and predetermine decision-making strategies would be too costly for animals to maintain. Similarly, commitment for extended periods of time to one strategy without the ability to adjust could be deleterious [23]. These authors suggested, with support from computational models, that temporal commitment to certain strategies with intermittent interruption to assess costs and switch strategies would be more advantageous for real world scenarios. Bumblebees in our study seem to follow a similar overall approach, as they first learn an abstract concept (relative size) and stick with this rule for approximately 20 bouts, at which point they change to a new strategy (see figure S2 for the individual difference between bees). A decrease in the number of choices taken to find all rewarding stimuli (figure 4) indicate that bees may have changed strategies to become more efficient. Further studies are needed to check the role of efficiency in strategy selection in animals. Further studies should involve videotaping the behaviour of bees during the training and test phases, so that one can make some direct inferences about time invested, mechanisms of inspecting stimuli and the efficiency of decisions.

In this light, our results support the idea that animals can adaptively weigh the costs of cognitive effort across decision-making approaches and choose the less cognitively demanding strategy [24]. This interpretation requires that the win-stay/lose-switch strategy was actually simpler than the relative size rule. Indeed, the win-stay/lose-switch heuristic is cognitively less demanding than any relational rule, simply because it is based only on the outcome of the previous choice, and therefore could be solved using working memory alone [25]. Accordingly, bees could have stored the visual template of the first stimulus in working memory and, if the first choice was correct, subsequently chosen a stimulus that had more overlap with the stored template, or if the first choice was incorrect, subsequently chosen a stimulus with less overlap (template hypothesis; [26]). The win-stay/lose-shift strategy has been broadly observed and explored in bees foraging strategies and flower constancy amongst variable rewarding species of flowers [27–31]. This type of sequential matching/non-matching to sample strategy has been shown to be solvable with a simple computational model based on the known neural circuitry of the bee brain, without requiring any higher-order abstract concept [6]. Learning and applying an abstract concept like relative size rule requires a substantial abstraction process to different stimuli that must work independent of the physical characteristics of stimuli [32]. In mammals, it is assumed that higher cognitive functions processed in the prefrontal cortex or analogous structures are essential for rule learning [33,34]. In insects, it has been proposed that rule learning occurs in the mushroom bodies, high-level sensory integration centres [35,36]. In contrast to rule-learning, bees can use a simple associative mechanism to remember the previously visited stimulus in order to make decisions about a subsequent stimulus. Therefore, the effort required in a win-stay/lose-switch type mechanism is likely to be lower than an abstract rule because bees can learn to recognize and associate a stimulus with reward without using their mushroom bodies [37,38]. For example, honeybees with inactivated mushroom bodies can perform some odour learning tasks as well as control bees [37,39]. Further, a realistic computational model of olfactory information processing in the bee brain shows that two parallel odour pathways with different functions provide the flexibility necessary for comparing multiple olfactory stimuli during associative and non-associative discrimination tasks [38].

Although our results indicate that bees switched to a win-stay/lose-switch heuristic, it is unclear why bees would learn the relative size concept first if the win-stay/lose-switch strategy is cognitively simpler. We speculate that this strategy may have been initially favoured simply to reduce the load on long-term memory and to speed up the decision-making process so as to avoid the quinine containing discs. During pretraining, bees only received reward from white disks. When training began, all of the discs suddenly contained coloured shapes and the bees found not only reward but also aversive quinine. Because of this abrupt and dramatic change, bees’ priority may have been to learn to avoid the quinine containing discs. To accomplish this quickly, they could have extracted a set of elementary visual features to avoid in the first bout of training. During the next bouts, instead of switching to a new strategy relying on working memory, they stuck with identifying and avoiding the template for the quinine containing discs. Over the next trials, they learned to generalise and group visual features across stimuli in a manner consistent with the concept of relative size [32,40]. Because constant monitoring of how well theywere doing would be too costly [23], it took them some time to assess their performance and try out a new strategy. Of course, further analysis of bees’ behaviour during the training and test phases are required to uncover the true mechanisms underlying bees’ strategy selections.

As a result of bees learning a relative size rule early in training, we would have expected to see an improvement on second choice performance from the first 10 bouts to the second 10 bouts in the training phase similar to the bees’ improvement on first choices (figure 2*b*). However, bees’ performance on second choices was not significantly different from chance level within 20 bouts of training. We are unable to say from our data why this was the case, but speculate that motivation and attention may play a role – once bees found reward, they might have been less likely to fly back within the arena to view stimuli head on to properly view and assess stimuli, and rather flew directly to a nearby disc to check for food, which statistically would be more likely to be unrewarding (because of the remaining five discs only two would be rewarding). This type of motivational-based exploration may also account for why bees eventually changed to a win-stay/lose-switch strategy. Figure S2 shows a large variability between individuals in second choice performance, and therefore individual differences in motivation and attention may have played a part in why second choice performance was lower than expected [41,42]. However, many of the bees did show an improvement in their second choices from the first 10 bouts to the second ten bouts. Analyses of sequential choices in future studies of animal cognition will help resolve these questions.

## Authors’ contributions

H.M. and L.C. conceived the study. H.M. designed and performed the experiment. H.M. and C.S. analysed data. H.M., C.S. and L.C. wrote the paper.

## Acknowledgments

We thank Stephan Wolf and Mark Roper for helpful discussions.

## Ethical statement

There are currently no international, national or institutional guidelines for the care and use of bumblebees in research. However, experimental design and procedures were guided by the 3Rs principles. Bumblebees were cared for on a daily basis by trained and competent staff, which included routine monitoring of welfare and provision of correct and adequate food during the experimental period.

## Supplemental information

Information includes two figures and one data file.

