## Supplemental figures for "Bumblebees learn a relational rule but switch to a win-stay/lose-switch heuristic after extensive training"

### Supplementary figure

(a)

| | Total area (cm²) | Edge length (cm) | Convex hull (cm²) |
|---|---|---|---|
| Small circle | 0.89 | 3.36 | 0.89 |
| Middle circle | 3.04 | 6.18 | 3.04 |
| Big circle | 6.46 | 9.01 | 6.46 |
| Small rectangle | 1.09 | 4.22 | 1.09 |
| Middle rectangle | 5.22 | 9.42 | 5.22 |
| Big rectangle | 9.06 | 12.48 | 9.06 |
| Small cross | 0.98 | 5.20 | 2.54 |
| Middle cross | 2.22 | 8.60 | 3.15 |
| Big cross | 4.50 | 11.44 | 6.42 |
| Small star | 0.56 | 5.00 | 1.10 |
| Big star | 3.45 | 12.30 | 6.74 |

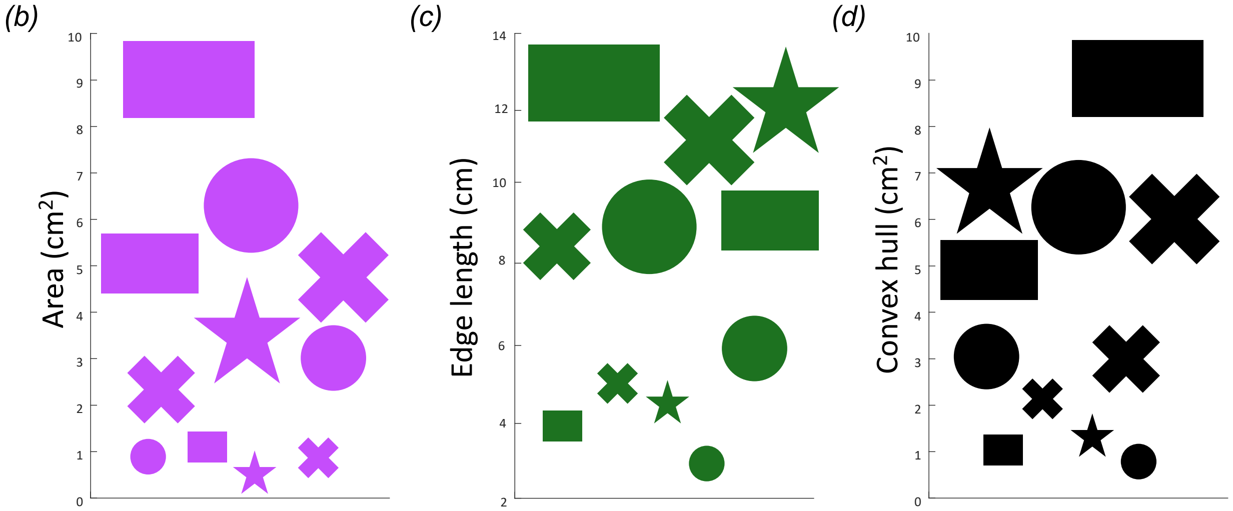


**Figure S1: Physical features of stimuli.** (*a*) Calculated total area, edge length and convex hull of each stimulus used in the study. (*b-d*) Plots visualizing variations of each stimulus feature across shapes. Note that for each feature several certain sized stimuli of one shape were closer in measure to a different size of another shape. For example, the big star was closer in total area to the middle circle and cross, while the middle rectangle was similar to the big circle and cross. This variation prevented bees from associating general sizes with reward and required the bees to compare relative sizes amongst shapes presented during each bout.

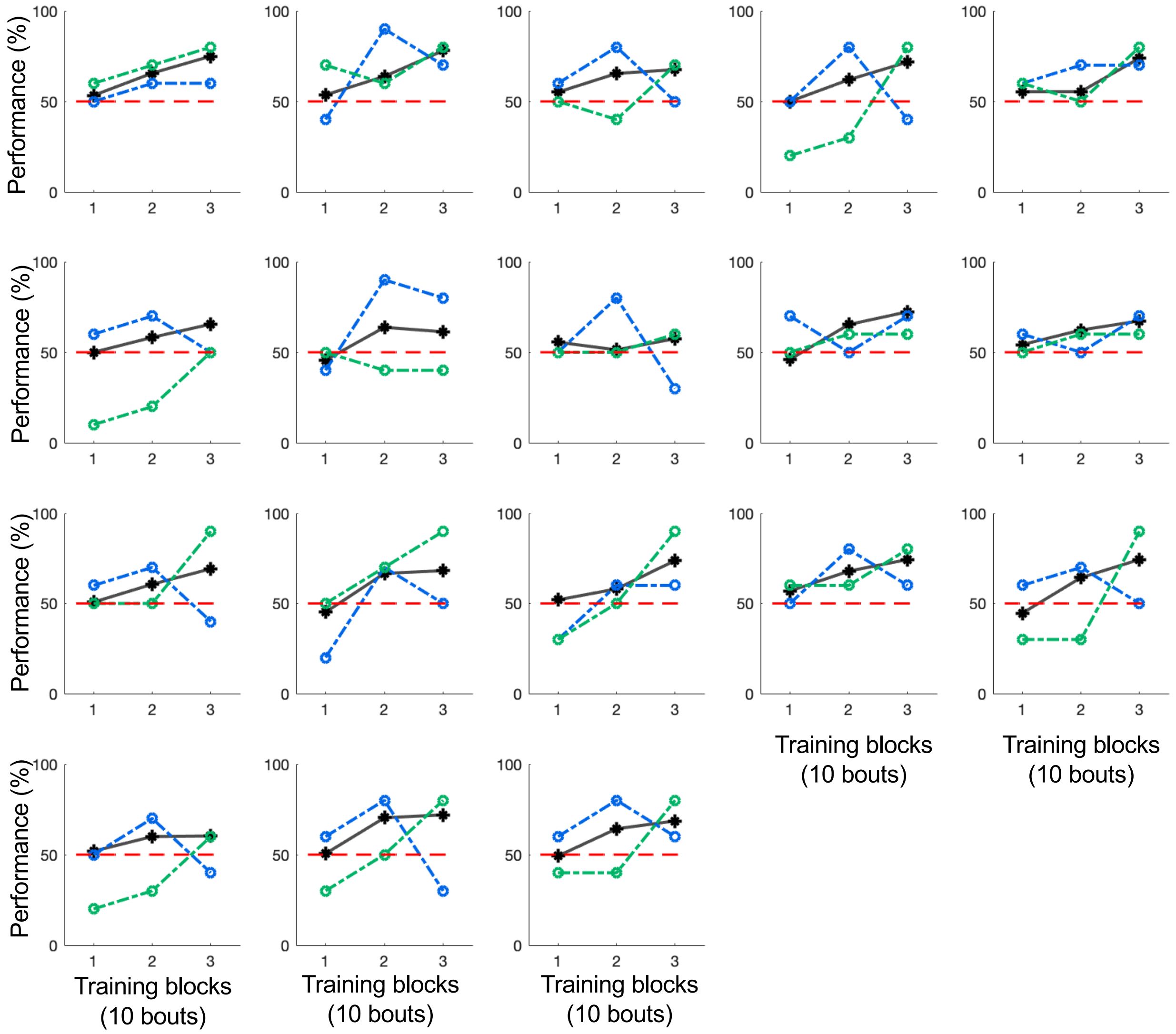


**Figure S2. Performances of individual bees in the training phase plotted for three different conditions.** Black lines show the learning curve of each bee based on the total choices. The blue and green curves display the bees' performances calculated from the first and second choices within each bout, respectively. Red dashed line = chance level (50%).
